# Direct pathway enrichment prediction from histopathological whole slide images and comparison with gene expression mediated models

**DOI:** 10.64898/2026.03.02.709137

**Authors:** Arfa Jabin, Shandar Ahmad

## Abstract

Molecular profiling of tumours via RNA sequencing (RNA-seq) enables clinically actionable stratification but remains costly, tissue-intensive, and time-consuming. Recent advances in computational pathology suggest that routine H&E whole-slide images (WSIs) can be utilized to estimate transcriptomic states of cancer cells. Given the WSI-derived predictions of transcriptional signatures are noisy, their use for accurate biological interpretation faces challenges. On the other hand pathway enrichment analysis has been routinely used in describing biologically meaningful cellular states from noisy gene expression data and some studies have evaluated the ability of WSI-predicted gene expression profiles to reconstruct enriched pathways in experiments where the two data modalities were concurrently available. However, it remains unclear if a predictive model that is designed to predict enriched pathways directly from WSI samples would be better than the current approaches to do so by first predicting gene expressions. Here, we develop and evaluate these two complementary approaches for predicting pathway enrichment profiles from WSIs in TCGA Breast Invasive Carcinoma (TCGA-BRCA) by training parallel models which predict pathway enrichment directly from image features and those which rely on predicted gene expression profiles, which is the current state-of-the-art. Our results suggest that under controlled experiments direct prediction of a selected pool of enriched pathways outperforms the models trained on predicting gene expression and then inferring enrichments on predicted gene expression values. These findings will be helpful in prioritizing the goals of predictive modeling of WSI images and improving diagnostic outcomes of cancer patients.

## 1. Introduction

Cancer is driven by genetic and regulatory alterations that reprogram cellular behaviour and reshape tissue architecture (Hanahan and Weinberg, 2011). Measuring tumour gene expression and pathway activity provides prognostic and therapeutic value, but RNA-seq and related assays can be resource-intensive and may delay treatment decisions (Wang et al., 2009). Histopathology remains the clinical gold standard for diagnosis, yet manual inspection is limited in its ability to resolve underlying molecular drivers and is inherently subjective (Madabhushi and Lee, 2016).

Deep learning has transformed digital pathology by learning reproducible representations from WSIs and associating morphology with molecular phenotypes, including mutations (Coudray et al., 2018), microsatellite instability (Kather et al., 2019), and transcriptomic profiles (Schmauch et al., 2020). However, most image-to-omics work emphasizes individual genes or single alterations. In contrast, biological pathways represent coordinated gene networks controlling oncogenic processes (e.g., cell cycle, apoptosis, PI3K/AKT, p53, MYC, Notch). It is well known in systems biology that pathway enrichment studies can be more robust, clinically interpretable, and actionable than single-gene outputs (Khatri et al., 2012; Subramanian et al., 2005). Initial evidence suggests that pathway activities can be inferred from histology images with attention-driven models, including in breast carcinoma where certain pathways are predicted with moderate-to-high discrimination (Qu et al., 2021). However best way to infer pathway enrichment from slides is still unclear.

This study proposes a WSI-based pipeline for predicting pathway activity using both **direct** and **indirect** strategies, aiming to bridge tissue morphology with transcriptome-derived functional states to support “virtual” molecular profiling from routine diagnostic slides.Direct approach aims to develop a multi-class prediction model, with enrichment of pathway in a sample being represented by corresponding binary values in the target vector. Indirect approach first trains the model to predict gene expression values and post-training converts predicted gene expressions to pathway enrichment. Using similar conditions, we find that direct pathway prediction is much more effective than its indirect counterpart, suggesting a novel strategy to interpret systems-level states of cancer tissue slides.

## 2. Methods

### 2.1 Study design and cohort construction (TCGA-BRCA)

We used TCGA Breast Invasive Carcinoma data (The Cancer Genome Atlas Network, 2012; Weinstein et al., 2013) accessed via the NCI Genomic Data Commons (GDC).

Cohort assembly consisted of 1231 available RNA-seq available profiles for primary tumour cases. Of these 1193 also had histology information. After QC and file processing a total of **987** cases with complete usability in which WSI and RNA-seq data were available and the same has been used in this study.

### 2.2 Whole-slide image preprocessing and tissue masking

Overall pipeline for image pre-processing is explained in Figure 1. In summary, WSIs were read using OpenSlide and standard image formats via PIL/Pillow (Goode et al., 2013). Images were converted between PIL and NumPy as needed. For each RGB image, tissue content was quantified as the proportion of non-masked pixels (non-background), enabling QC and filtering of low-tissue images. Following **filtering operations (artifact suppression + enhancement) were implemented:**

**Figure 1.**
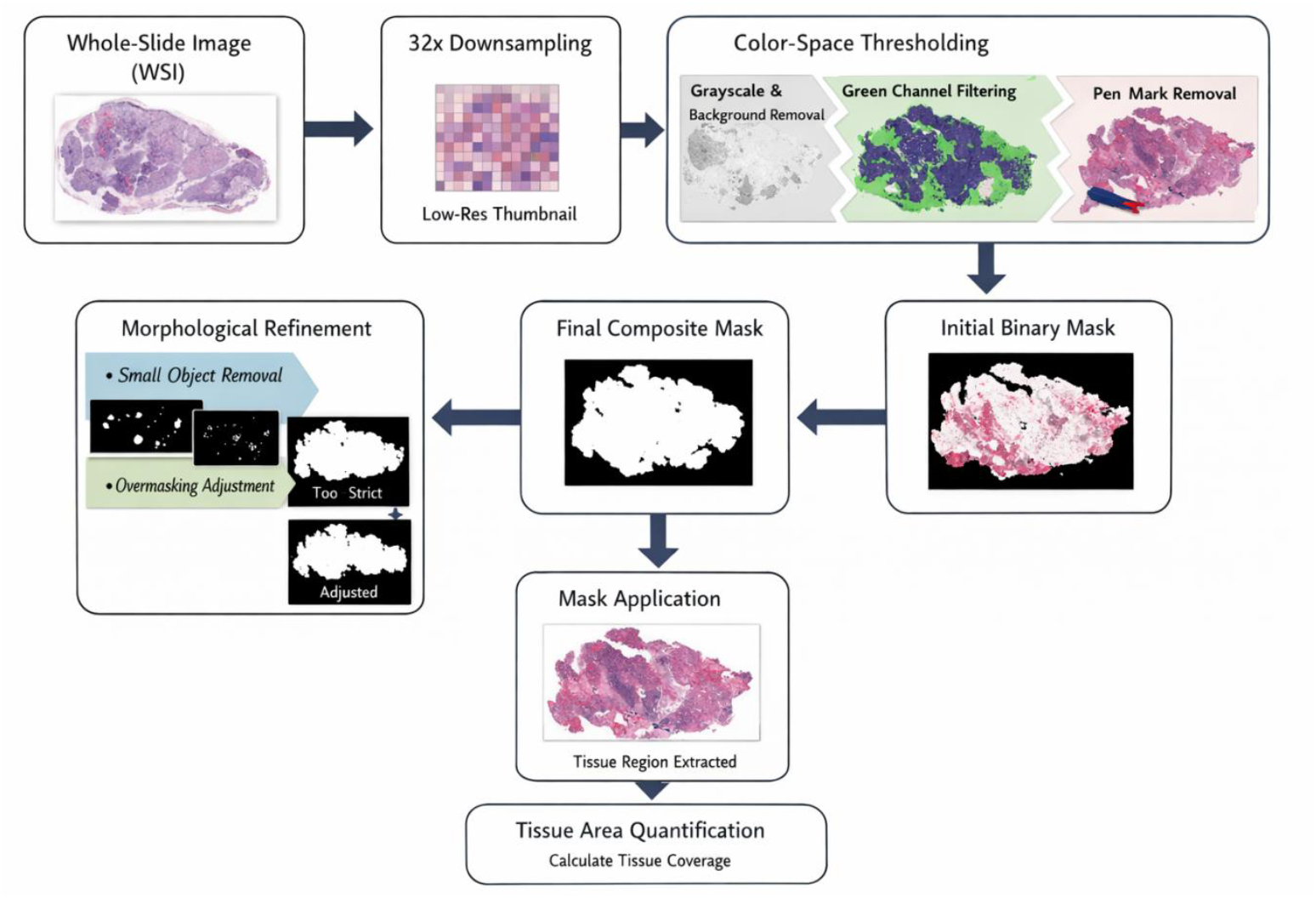
Automated histopathology image prefiltering pipeline.

1. **Colour/intensity transforms:** RGB images were converted to grayscale and/or HED (hematoxylin/eosin/DAB) via colour deconvolution principles (Ruifrok and Johnston, 2001).
2. **Contrast enhancement:** Histogram equalization / adaptive histogram equalization (CLAHE-style variants) to improve local contrast (Pizer et al., 1987; Zuiderveld, 1994).
3. **Otsu thresholding:** Global tissue/background segmentation using Otsu’s method (Otsu, 1979).
4. **Morphological refinement:** Erosion/dilation/opening/closing, hole filling, removal of small objects/holes to denoise masks and preserve contiguous tissue. Implemented using scikit-image and SciPy (van der Walt et al., 2014).
5. **Colour-based artifact filters:** Removal of grays/non-tissue colours (background/glass) was performed by sequential use of Green channel suppression, specific pen mark removal (red/green/blue pen filters via channel logic), near-black and near-white removal (edges/background) steps.
6. **Recursive “safety loop” QC:** If masking exceeded pre-set thresholds (over-masking risk), hyperparameters were automatically adjusted and filtering repeated to retain informative tissue.

### 2.3 Deep feature extraction from WSIs

Tissue image tiling (partitioning of images into rectangular tiles) was first carried out by creating non-overlapping 224 × 224 pixel patches extracted from segmented tissue (CNN-native resolution). Tiles with Tissue threshold < 20% tissue area were discarded. Finally, if an image has more than 8,000 tiles satisfying these conditions, a maximum of 8000 were retained for training a multi-instance model. The process is given in more detail in Table 1.

**Table 1.** Tissue coverage metrics for preprocessing: (A) QC. Tissue %: valid pixels corresponding to tissue (pink/purple/brown). *Target:* high enough to cover the specimen. Masked %: invalid pixels (glass/background). *Target:* sufficient to remove background without being excessive (B) WSI preprocessing and feature extraction hyperparameters

| Metric | What it Represents | Ideal Value |
| --- | --- | --- |
| Tissue % | Valid pixels (Pink / Purple / Brown tissue regions) | High enough to capture the entire tissue sample |
| Masked % | Invalid pixels (White glass / Black background) | High enough to remove all background, but not excessive |

(B)
| Parameter | Value | Description |
| --- | --- | --- |
| Tile Size | 224×224 | Spatial dimensions of extracted patches |
| Backbone | ResNet50 | Pre-trained on ImageNet, truncated at GAP |
| Max Tiles (N) | 8000 | Maximum number of patches extracted per WSI |
| Tissue Threshold | 20% | Minimum tissue area required to keep a tile |
| Output Dim | 2048 | Dimensionality of the resulting feature vectors |

### 2.4 Feature encoding (ResNet50)

Tile embeddings for each tile were computed using a ResNet50 CNN pretrained on ImageNet (Deng et al., 2009; He et al., 2016). Final classification head was removed and features were extracted at the Global Average Pooling (GAP) layer. This resulted in 2048-dimensional embedding per tile or N × 2048 bag (up to 8,000 × 2,048) pool of image representation for N tiles. Feature aggregation was carried out as in Figure 2.

**Figure 2.**
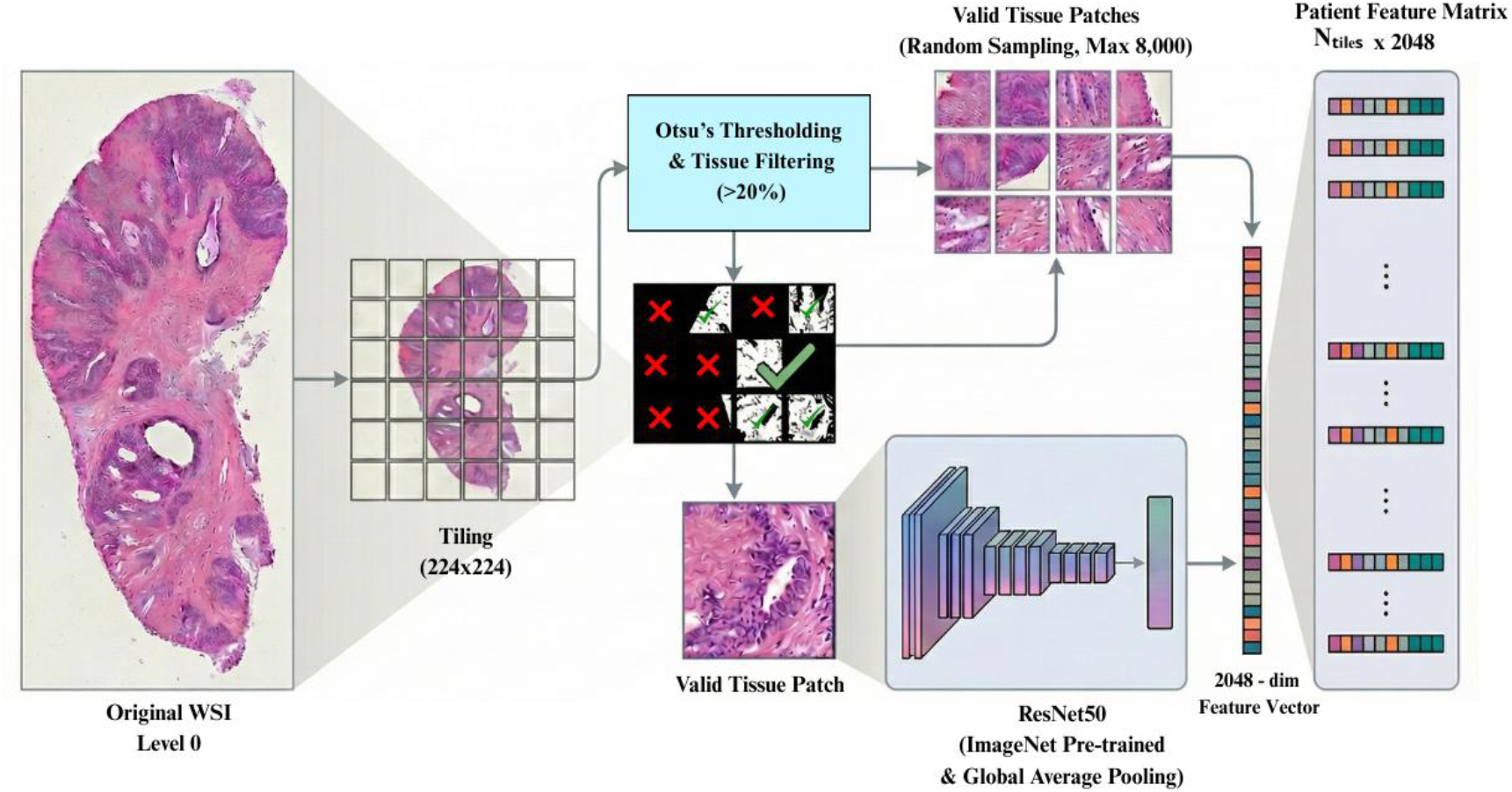
WSI processing and feature extraction pipeline.

### 2.5 RNA-seq quantification and normalization

RNA-seq was aligned with STAR (Dobin et al., 2013). We used FPKM-UQ (upper-quartile normalized FPKM) as the expression measure (GDC workflow convention; TCGA/GDC documentation). Expression tables were filtered to valid Ensembl IDs and technical artifacts removed. For each of the 987 patients, expression across 60,660 Ensembl loci was Z-scored within-sample (mean 0, variance 1), emphasizing relative gene ranking per tumour. The top 100 genes by absolute Z-score were extracted as a patient-specific dominant transcriptional signature.

### 2.6 Pathway activation profiling and filtering

Pathway inference used KEGG 2021 Human as the reference pathway database (Kanehisa & Goto, 2000). Ensembl IDs were mapped to HGNC symbols prior to enrichment. Over-representation analysis (ORA) was used to identify enriched pathways per patient (methodologically aligned with standard pathway analysis practice Khatri et al., 2012, Subramanian et al., 2005). Gene status as “Active/Inactive” for binarization was made by carrying out a pathway enrichment analysis and different thresholds which included p<0.05, 0.01, 0.001, 0.0001. First top 20 enriched pathways are collected from each patients and then a pool from all such sets was created. Across the cohort Frequency near-ubiquitous pathways (>90% prevalence; e.g., Ribosome) and rare pathways (<10% prevalence) were removed from the list to avoid sparsity. The remaining pooled set consists of 40 pathways, leading to tge status matrix of 987 × 40 (samples × pathways), setting up the data for the multi-class prediction model.

### 2.7 Modeling approaches

In general WSIs is first downsampled and converted to NumPy. Parallel filtering then removes background (green/gray suppression) and artifacts (pen filters), with a recursive safety loop preventing over-masking. (3) Binary morphology refines the tissue mask (small object removal, hole filling). (4) Output: clean masked tissue image plus tissue/background coverage metrics

Overall pipeline for pathway prediction in GE-mediated and direct models is presented in Figure 3 and Figure 4 respectively. The same are explained below.

**Figure 3.**
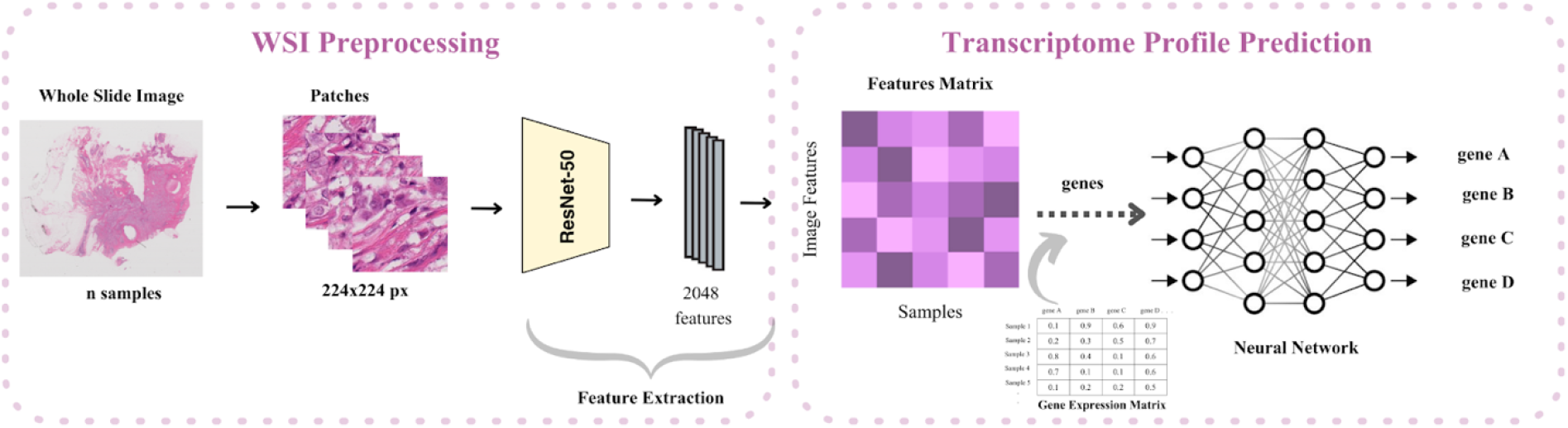
Workflow to predict pathways of tumours from whole slide images based on gene expression prediction (Indirect)

**Figure 4.**
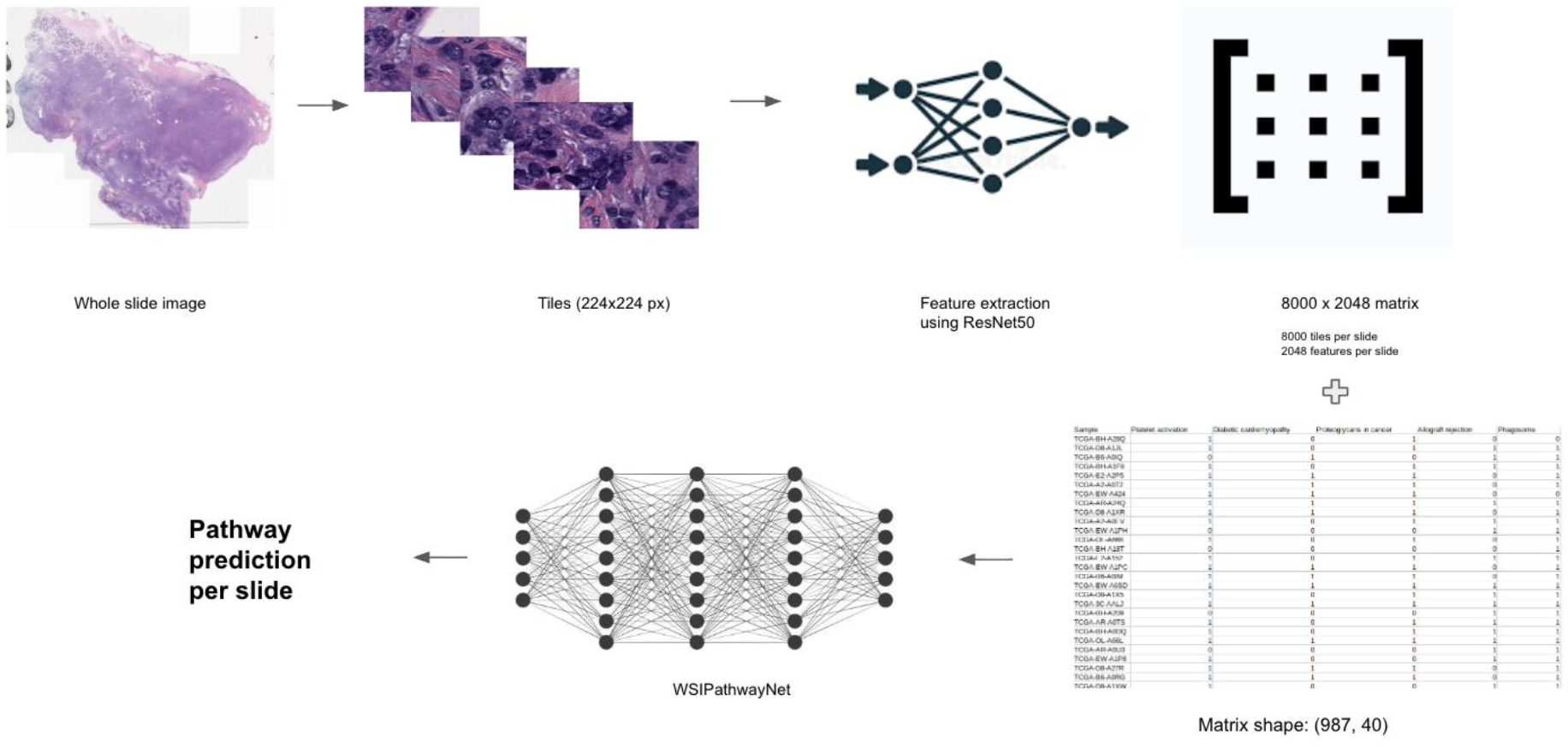
Workflow to predict pathways of tumours from whole slide images (Direct)

**Figure 5.**
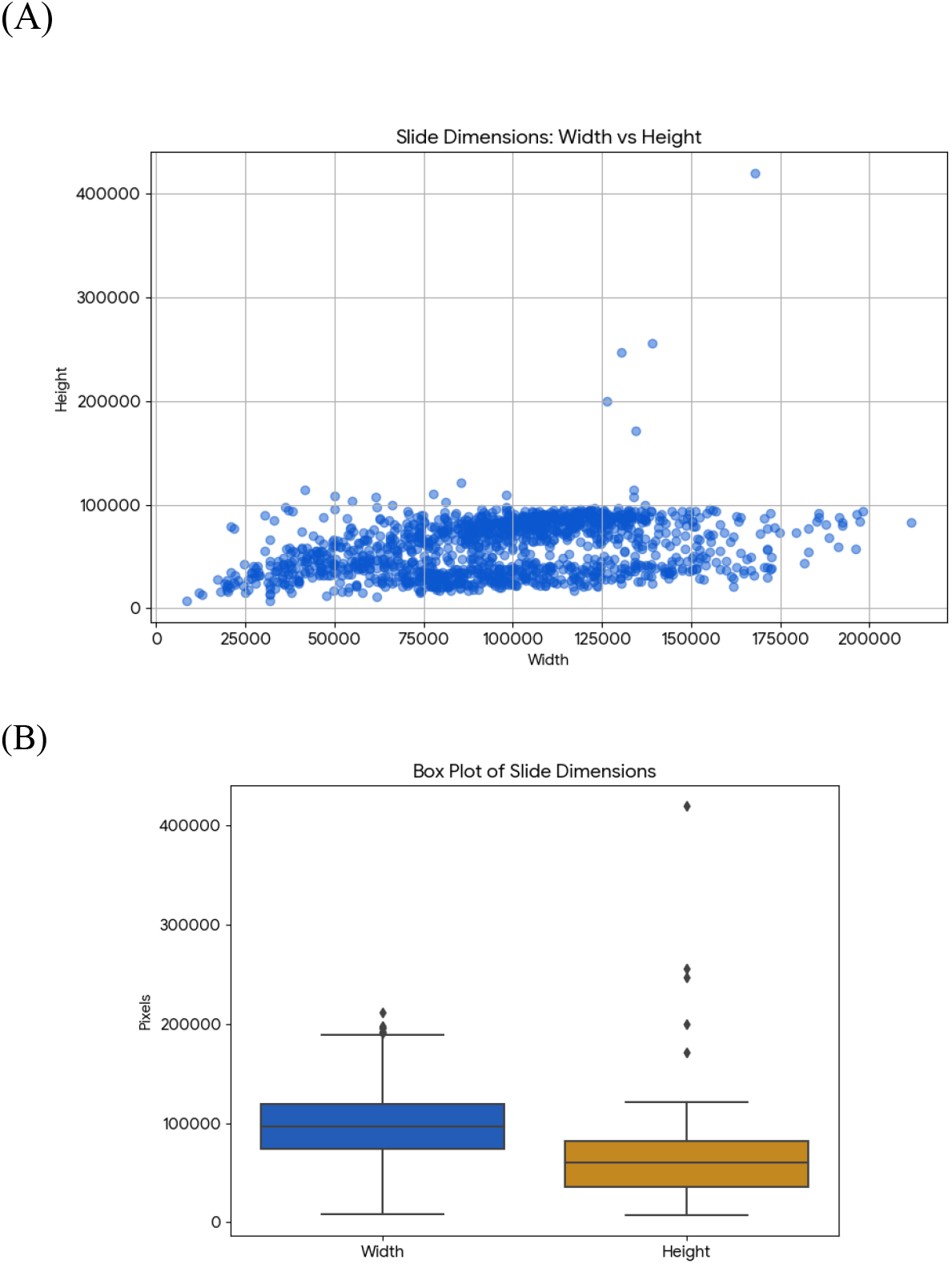
Slide dimension heterogeneity: (A) Scatter plot of slide width vs. height (pixels) for N=1,434 WSIs, showing broad variance without a single standardized size. (B) Box plots of width and height distributions highlighting outliers, including an extreme height

#### 2.7.1 Prediction of enriched pathways using Gene expression mediated (indirect) model

This approach consisted of two steps:

##### Stage I: Gene expression prediction from WSIs

Slide-level ResNet-derived embeddings were used to predict gene expression through supervised regression (Jabin et al., 2025). A multi-layer perceptron (MLP) with SiLU activations (Ramachandran et al., 2017), Layer Normalization (Ba et al., 2016), and dropout regularization (Srivastava et al., 2014) was employed. The final layer consisted of a linear regression head producing continuous gene expression estimates. Optimization was performed using Adam (Kingma and Ba, 2015) with mean squared error loss. To manage the large transcriptomic target space, training was conducted in gene batches with checkpointing to ensure computational efficiency and stability.

##### Stage II: Pathway activity from predicted transcriptome

Using predefined pathway gene sets (40 KEGG pathways), predicted gene expression values were aggregated into pathway activity scores. Continuous pathway scores were retained for regression evaluation; then binarized for classification.

#### 2.7.2 Direct prediction of enriched pathways (direct prediction) model

This approach is similar to GE prediction in the indirect approach. However, the model is now trained to predict 40-dimensional binary classes instead of regression predictions used for gene expression prediction.

#### 2.7.3 Regression metrics and Training protocol

Pearson correlation, Spearman correlation, MSE and Matthews correlation coefficient MCC) are used to assess model performances where applicable. The dataset was partitioned using a stratified splitting strategy to preserve class distribution across training, validation, and test sets. Model optimization was performed using the AdamW optimizer with decoupled weight decay regularization (Loshchilov and Hutter, 2019). A ReduceLROnPlateau scheduler was employed to adaptively lower the learning rate when validation performance plateaued. To mitigate overfitting, early stopping was applied based on validation AUC, terminating training when no further improvement was observed over a predefined patience window. To ensure robustness to random initialization and data partition variability, models were trained across multiple independent random seeds. Final pathway probabilities were obtained by aggregating predictions across seeds and splits, yielding a consensus global prediction matrix for downstream evaluation.

### 2.8 Architecture of training model “PathwayNet”

Pathway activation was modeled using a supervised multi-layer perceptron (MLP) classifier operating on slide-level feature embeddings. The network received fixed-dimensional representations derived from whole-slide image aggregation as input. These features were processed through a compact hidden layer with nonlinear activation (SiLU) to enable smooth gradient propagation and expressive feature transformation. To improve generalization and reduce overfitting, normalization and dropout regularization were applied within the hidden layers. The output layer employed a sigmoid activation function to estimate the probability of pathway activation. The model was trained independently for each pathway target, and the resulting predictions were assembled into a consolidated slide-by-pathway probability matrix aligned with the ground-truth pathway annotation matrix.

#### 2.8.1 Class imbalance handling

To address pathway sparsity, we used SMOTE oversampling, with neighbour count dynamically adjusted to the minority class size (Chawla et al., 2002).

## 3. Results

### 3.1. Image size and usable tile frequencies in the data set

Figure 1 shows the distribution of image sizes across studies curated for this study. It is visible that the patient slides have a varying width, although heights were consistent across batches pooled here. Even though image sizes are different, tiling of the image can reduce the biased representation in the final predictive model.

### 3.2 Direct and indirect prediction of pathway enrichment from WSI data

Figure 6 shows the prediction performance of each model when pathway enrichment is predicted directly from slide features, Figure 7 shows the MCC between predicted and ground truth values for the same models. The bar chart shows that the prediction scores are robust ranging from MCC=0.3 and up to 1.0 for some pathways. Mean AUROC was found to be 0.931 and mean MCC reached 0.7291. In contrast MCC from the GE-mediated model was much smaller i.e. just 0.64. Thus we observe a substantial gain in MCC scores for models trained on pathway enrichment directly. Additional prediction performance scores for GE-mediated prediction models are shown in Table 2.

**Figure 6.**
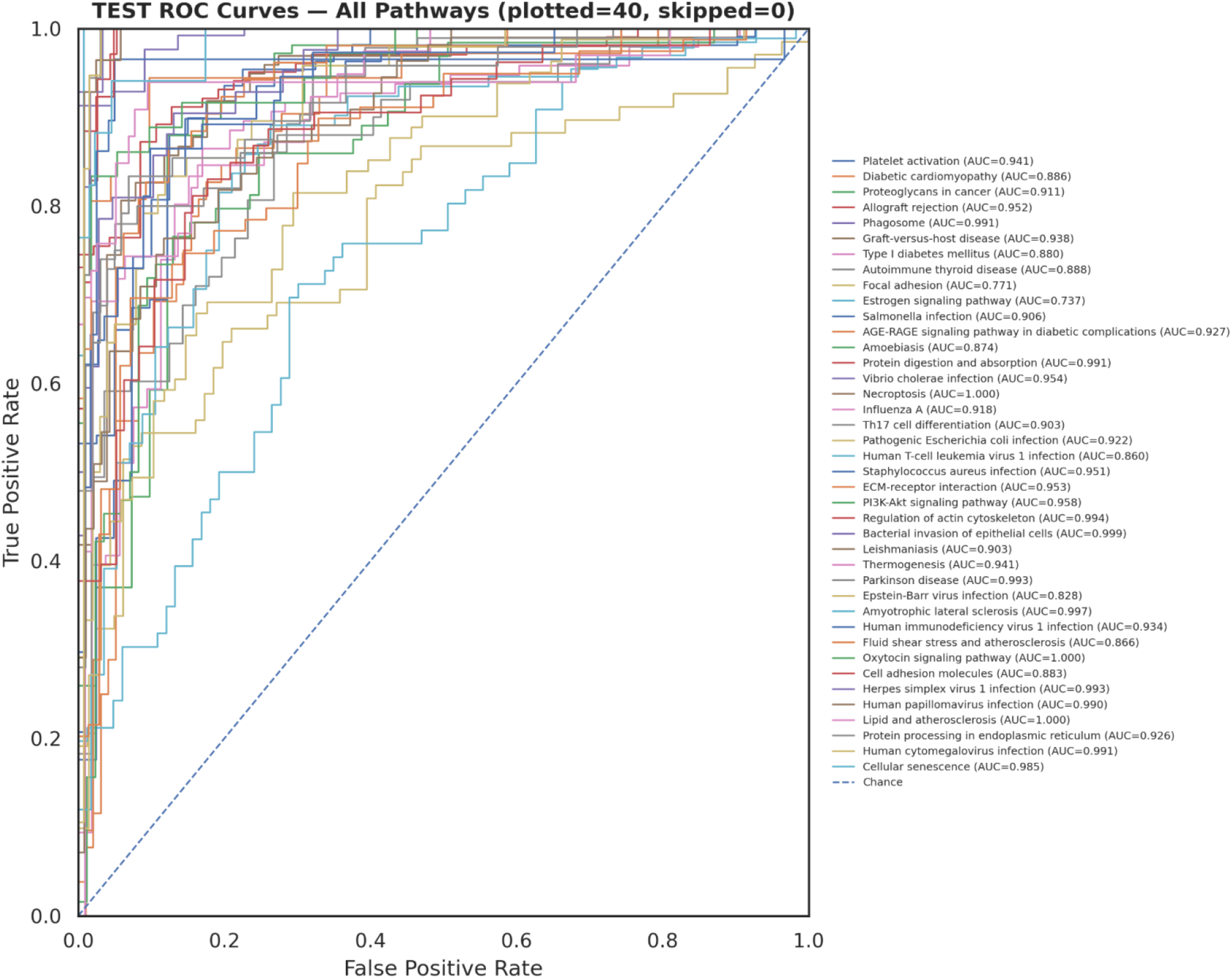
Binary-class prediction ROC curves for different pathways included in the target vector using direct pathway prediction method (without GE prediction).

**Figure 7.**
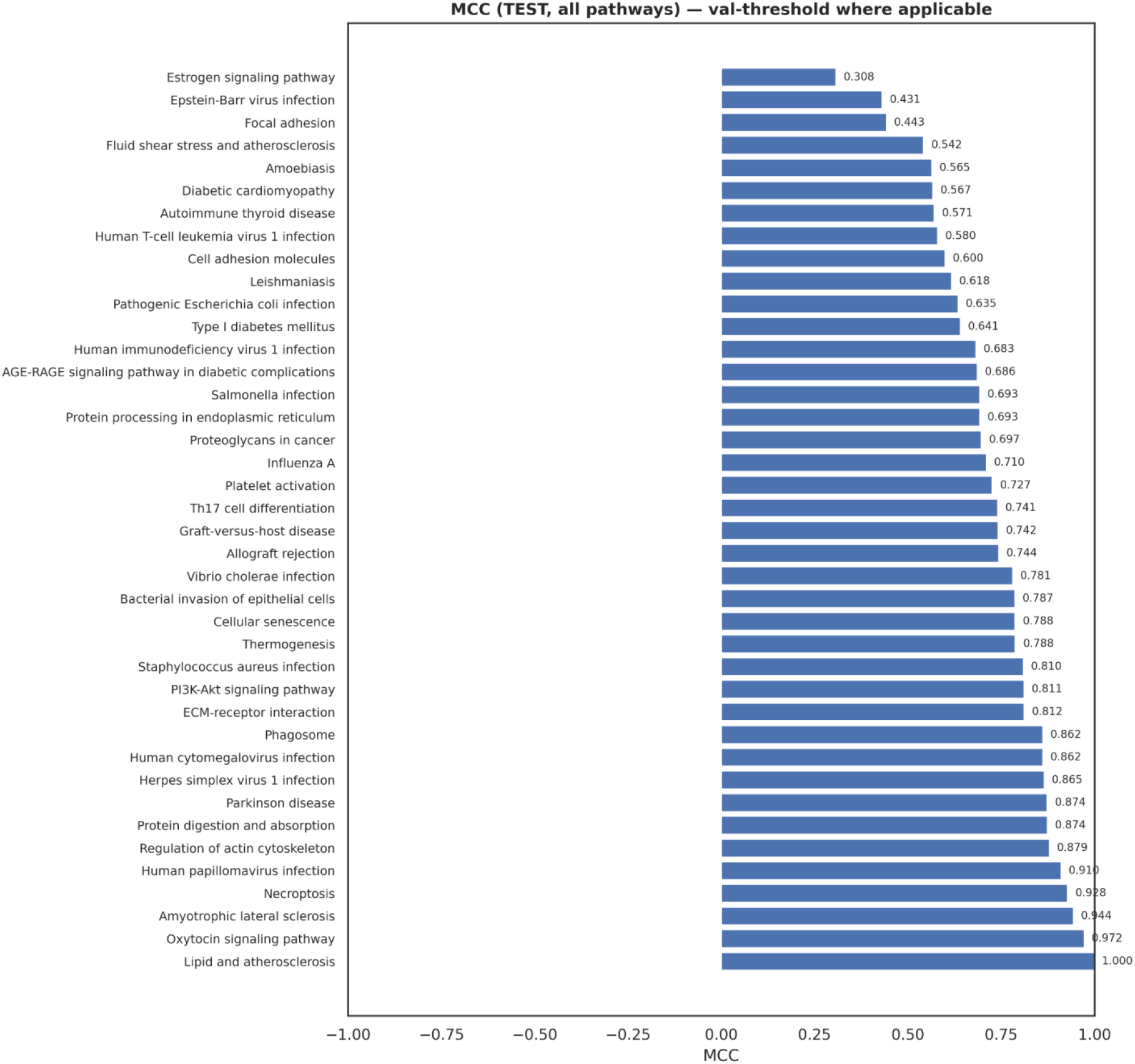
AUC of ROC for the pathway binary class prediction for each of the selected pathway using direct pathway prediction method (without GE prediction).

**Table 2.** Indirect framework classification performance across significance thresholds.

| p-value<br>Threshold | Accuracy | Precision | Recall | F1-Score | p-value<br>Threshold |
| --- | --- | --- | --- | --- | --- |
| 0.05 | 62.8% | 0.434 | 0.087 | 0.145 | 0.05 |
| 0.01 | 63.8% | 0.509 | 0.047 | 0.086 | 0.01 |
| 0.001 | 64.2% | 0.638 | 0.025 | 0.048 | 0.001 |
| 0.0001 | 64.0% | 0.670 | 0.014 | 0.028 | 0.0001 |

## 4. Discussion

Highest-performing pathways were enriched for immune/inflammatory and microenvironment/ECM programs, which produce strong and spatially explicit morphology (e.g., lymphocyte infiltration, stromal remodeling) that is readily captured in H&E. In contrast, hormone signaling pathways (e.g., estrogen signaling) showed lower predictability, consistent with more intracellular signalling effects that may not map cleanly to overt tissue architecture.

These observations align with prior evidence that some pathway states are predictable from breast WSIs using deep learning and with broader computational pathology findings that tumour microenvironment signals are morphologically salient (Qu et al., 2021). This work compares two strategies for pathway inference from WSIs. The direct approach provides stronger classification performance, achieving mean AUROC near 0.93. The indirect approach follows a mechanistically aligned hierarchy (morphology to transcriptome to pathways), enabling gene-level interpretation and supporting biological explanations of pathway activation.

A key implication is that pathway-level prediction performance is driven by how directly a pathway’s biology manifests in histomorphology. Microenvironmental processes yield strong signal, whereas pathways dominated by intracellular signalling may require additional modalities or more targeted weak supervision to achieve comparable performance. This direction is consistent with multimodal frameworks that explicitly model pathway tokens and histology patch tokens (Jaume et al., 2024).

## 5. Conclusion

We developed a WSI-based framework for predicting pathway activity in TCGA-BRCA and empirically compared direct and indirect paradigms. The **direct model** achieved slightly higher pathway classification performance (mean AUROC 0.931; mean MCC 0.729). The **indirect model** provided biologically meaningful structure and gene-level interpretability, improving mechanistic understanding of predicted pathway states. Together, these results support the feasibility of “virtual molecular profiling” from routine histology and highlight tumour microenvironment biology as strongly encoded in H&E morphology.

